# Dissociable influences of reward and punishment on adaptive cognitive control

**DOI:** 10.1101/2020.09.11.294157

**Authors:** Xiamin Leng, Debbie Yee, Harrison Ritz, Amitai Shenhav

**Author notes:** Correspondence should be addressed to Xiamin Leng.

## Abstract

To invest effort into any cognitive task, people must be sufficiently motivated. Whereas prior research has focused primarily on how the cognitive control required to complete these tasks is motivated by the potential rewards for success, it is also known that control investment can be equally motivated by the potential negative consequence for failure. Previous theoretical and experimental work has yet to examine how positive and negative incentives differentially influence the manner and intensity with which people allocate control. Here, we develop and test a normative model of control allocation under conditions of varying positive and negative performance incentives. Our model predicts, and our empirical findings confirm, that rewards for success and punishment for failure should differentially influence adjustments to the evidence accumulation rate versus response threshold, respectively. This dissociation further enabled us to infer how motivated a given person was by the consequences of success versus failure.

**Author Summary:** From the school to the workplace, whether someone achieves their goals is determined largely by the mental effort they invest in their tasks. Recent work has demonstrated both why and how people adjust the amount of effort they invest in response to variability in the rewards expected for achieving that goal. However, in the real world, we are motivated both by the positive outcomes our efforts can achieve (e.g., praise) *and* the negative outcomes they can avoid (e.g., rejection), and these two types of incentives can motivate adjustments not only in the amount of effort we invest but also the *types* of effort we invest (e.g., whether to prioritize performing the task *efficiently* or *cautiously*). Using a combination of computational modeling and a novel task that measures voluntary effort allocation under varying incentive conditions, we show that people should and do engage dissociable forms of mental effort in response to positive versus negative incentives. With increasing rewards for achieving their goal, they prioritize efficient performance, whereas with increasing penalties for failure they prioritize performing cautious performance. We further show that these dissociable strategies enable us to infer how motivated a given person was based on the positive consequences of success relative to the negative consequences of failure.

## Introduction

People must regularly decide how much mental effort to invest in a task, and for how long. When doing so, they weigh the costs of exerting this effort against the potential benefits that would accrue as a result [1,2]. These benefits include not only the positive consequences of success (e.g., money or praise) but also the negative consequences of failure (e.g., criticism or rejection). Prior work suggests that people likely vary in the extent they are motivated by the prospect of achieving a positive outcome versus avoiding a negative outcome [3,4]. For example, some students study diligently to earn praise from their parents while others do so to avoid embarrassment. The overall salience of these incentives will determine when and how a given person decides to invest mental effort (i.e., engage relevant cognitive control processes [5], including when they choose to disengage from effortful tasks [6,7]). However, while a great deal is known about how people adjust cognitive control in response to varying levels of potential reward [5,8,9], much less is known about how they similarly adjust to varying levels of potential punishment, nor the types of control allocation strategies that are most adaptive under these two incentive conditions.

Previous research has examined how control allocation varies as a function of the reward for performing well on a task, such that participants generally perform better when offered a greater reward [10–14]. For instance, when earning rewards during a cognitive control task (e.g., Stroop) is contingent on both speed and accuracy, participants are faster and/or more accurate as potential rewards increase [11,15–17]. While studies have examined how motivation to avoid negative outcomes influence cognitive control [18–22], a challenge of interpreting these mixed behavioral patterns is that participants deploy a variety of behavioral strategies as potential punishments increase [22,23]. Past work has demonstrated that these strategies, such as increased task processing (e.g., attentional focus) or adjusting decision thresholds, can be linked to different forms of control adjustment (e.g., prioritizing speed versus accuracy; [24– 27]). However, it remains unknown whether participants selectively deploy different forms of control adjustments when incentivized under distinct incentive regimes (i.e., to avoid poor performance versus achieve good performance).

Recent theoretical work helps to frame predictions regarding when and how people might vary their control allocation in response to different forms of incentives [1]. For instance, normative accounts of physical effort allocation have proposed that animals and humans vary the intensity of their effort (e.g., motor vigor) to maximize their net reward per unit time (reward rate [28–31]). We have recently extended this framework to describe how people determine the appropriate allocation of *cognitive control* in a given situation. Specifically, we have suggested that people select the amount and type(s) of cognitive control that maximize the overall rate of expected rewards, while minimizing expected effort costs. The difference between these two quantities, referred to as the Expected Value of Control (EVC), indexes the extent to which the benefits of control outweigh its costs [1,2,32] (see also [33]).

The EVC model has been successful at accounting for how people vary the intensity of a particular type of control (e.g., attention to a target stimulus/feature) to achieve greater rewards [34,35]. However, limitations in existing data have prevented EVC from addressing how the *type* of control being allocated should depend on the type of incentive being varied. One limitation, noted above, is the dearth of research on how people adjust control to positive versus negative incentives. A second potential limitation is that most existing studies examine how performance varies over a fixed set of trials (e.g., 200 total trials completed over the course of an experiment). The maximal expected reward is determined by the number of trials in the task, which could limit the underlying drive to maximize reward rate. A stronger test of reward rate maximization, and one that is arguably more analogous to real-world effort allocation, would allow participants to perform as much or as little of the task as they like over a fixed duration [36], to tighten the link between reward rate and overall expected reward.

In the current study, we developed a novel paradigm in which participants perform consecutive trials of a control-demanding task (the Stroop task) over a fixed time interval. We examined how the amount and type(s) of control allocated to this task varied under different incentive types (reward vs. punishment) and different magnitudes of those incentives (small vs. large). Across two experiments, participants demonstrated distinct patterns of task performance in the two incentive conditions: faster responses for increasing rewards, slower but more accurate responses for increasing punishment. We show that these patterns are consistent with normative predictions of a control allocation model that maximizes reward rate while minimizing effort costs. The model predicts that rewards versus punishments favor divergent control strategies: higher reward promotes faster information processing to maximize (correct) response rate, whereas higher punishment promotes greater caution to minimize potential errors. Within the framework of a drift diffusion model (DDM), our normative model predicts that participants will respond to increases in reward level by both increasing their evidence accumulation rate (drift rate) and lowering their response threshold, whereas they will respond to increases in punishment level by primarily increasing their threshold. Model fits to behavioral data across both studies confirmed these predictions.

Our model’s ability to make divergent predictions about the influence of incentives on the *joint* allocation of two forms of control (i.e., across drift rate and threshold) enabled us to make further inferences based on each participant’s unique behavioral profile. Specifically, by estimating how these DDM parameters varied together across conditions, we were able to infer how sensitive that participant might have been to reward and punishment to generate the pattern of behavior that they did. Collectively, this work demonstrates a compelling novel method for inferring variability in how people evaluate costs and benefits when deciding when and how much to allocate cognitive control.

## Results

Participants (N=32) performed a task in which they were given fixed time intervals (between 8 and 12 seconds long) to perform as many trials as they wanted of a cognitively demanding task (Stroop task; Figure 1). They received monetary reward for each correct response within a given interval, and incurred a monetary loss (penalty) for each incorrect response. The magnitude of reward and penalty ($0.01 or $0.10) were independently varied across intervals, and were cued prior to the start of each interval.

**Figure 1.**
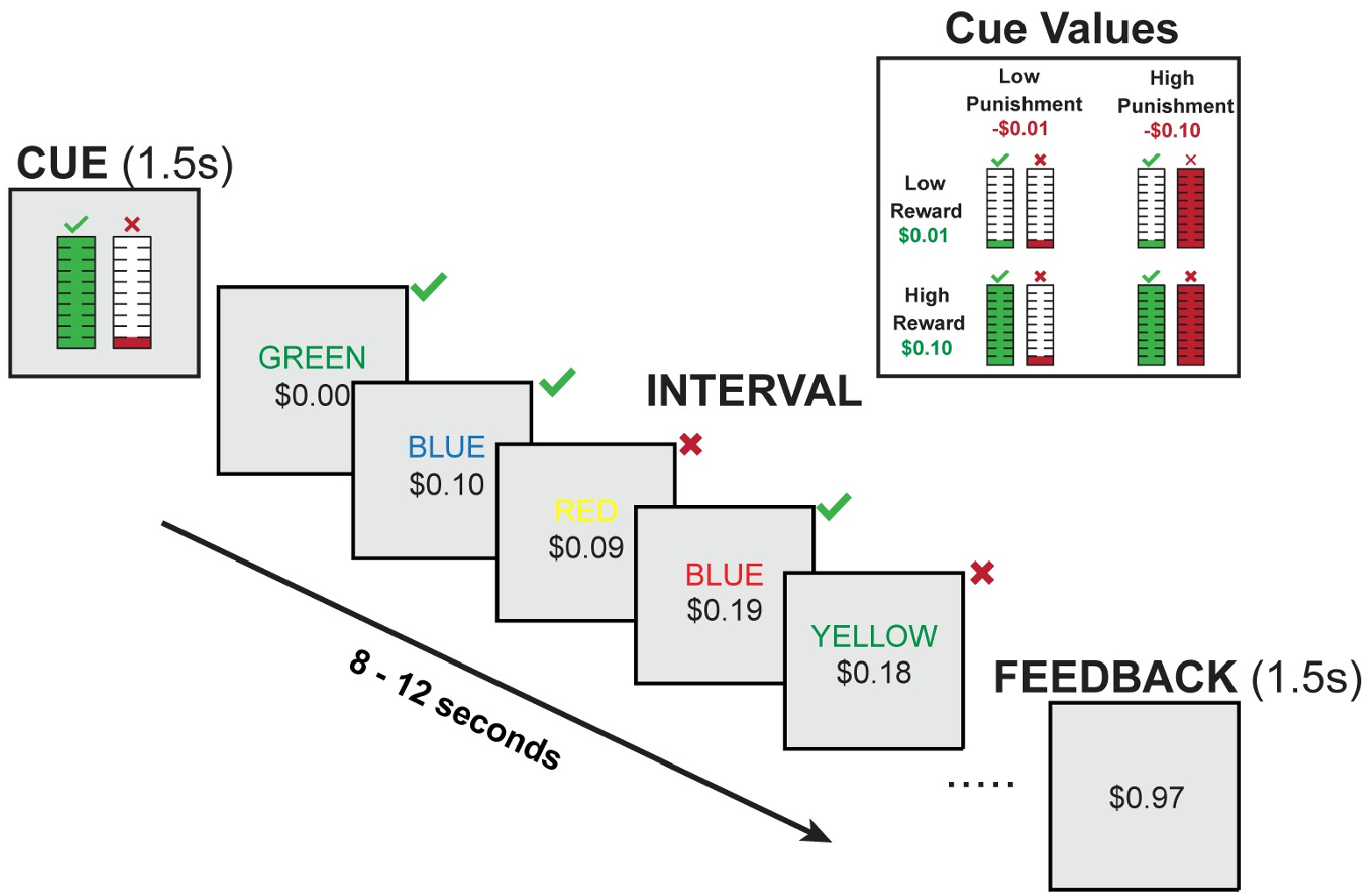
Interval-Based Incentivized Cognitive Control Task. At the start of each interval, a visual cue indicates the amount of reward (monetary gain) for correct responses and the penalty amount (monetary loss) for incorrect responses within that interval. Participants can complete as many Stroop trials as they want within that interval. The cumulative reward over a given interval is tracked at the bottom of the screen. Correct responses increase this value, while incorrect responses decrease this value. At the end of each interval, participants are told how much they earned. The upper right inset shows the cues across the four conditions.

### Behavioral Performance

We found that when participants were expecting a larger reward for each correct response, they completed more trials correctly in a given interval compared to when they were expecting smaller rewards (*F*_(1,31)_=28.72, *p*<0.001; Figure 2A, Table 1). Variability in punishment magnitude appeared to have the opposite influence on behavior. When participants were expecting a larger punishment for each incorrect response, they completed fewer correct trials in a given interval than when they were expecting smaller punishments (*F*_(1,31)_=23.11, *p*<0.001; Figure 2B). We also observed a trending interaction between reward and punishment (*F*_(1,29)_=3.77, *p*=0.062) whereby the reward-related improvements in interval-level performance were enhanced in high-punishment compared to low-punishment intervals.

**Table 1.** Mixed Model Results for Correct Responses per Second *: *p*<0.05, **: *p*<0.01, ***: *p*<0.001

| Correct Responses Per Second |  |  |  |
| --- | --- | --- | --- |
| *Predictors* | *Estimates* | *S.E.* | *P-Value* |
| Age | -0.036 | 0.031 | 0.238 |
| Female - Male | 0.075 | 0.032 | **0.020\*** |
| High Penalty - Low Penalty | -0.026 | 0.005 | **<0.001\*\*\*** |
| High Reward - Low Reward | 0.038 | 0.007 | **<0.001\*\*\*** |
| Average Congruence | -0.015 | 0.005 | **0.001\*\*** |
| Reward × Penalty | -0.009 | 0.005 | 0.052 |
| Number of Subjects | 32 |  |  |
| Observations | 2469 |  |  |
| Marginal $R^2$ / Conditional $R^2$ | 0.093 / 0.551 | | |

**Figure 2.**
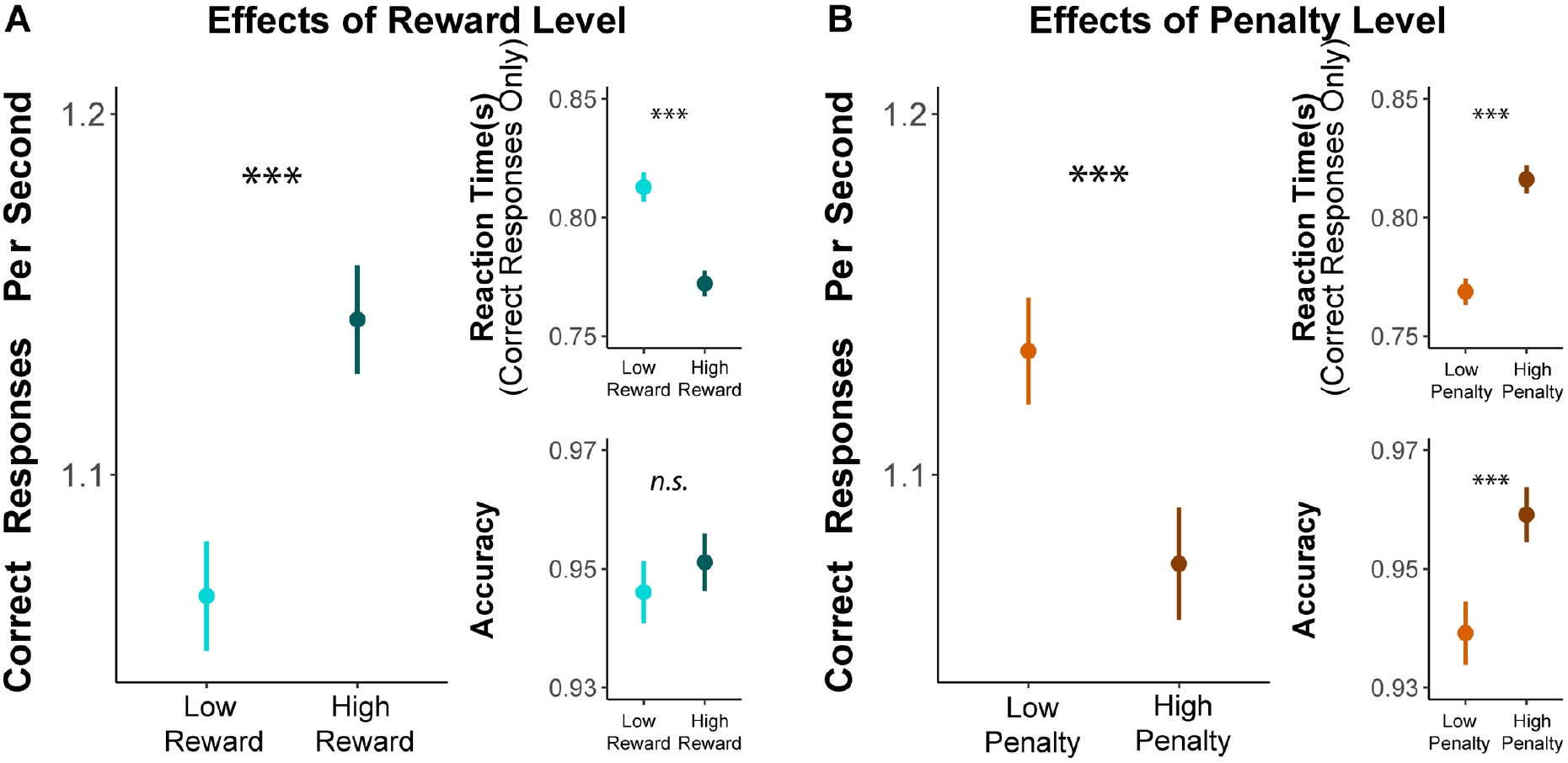
Effects of reward and punishment on overall task performance. **A)** With increasing expected reward, participants completed more correct responses per second within a given interval (**left**), which reflect faster responding on correct trials (**top right**) without any change in overall accuracy (**bottom right**). **B)** With increasing expected punishment, participants instead completed fewer trials per second over an interval, reflecting slower and more accurate responses. Error bars reflect 95% CI. n.s.: p>0.05; ***: p<0.001

When separately examining how incentives influenced speed and accuracy, we found an intriguing dissociation that helped account for the inverse effects of reward and punishment on the number of correct responses per second. We found that larger potential rewards induced responses that were faster (*F*_(1,28)_=31.83, *p*<0.001) but not more or less accurate (*Chisq*_(1)_=0.26, *p*=0.612; Figure 2A, Table 2). By contrast, larger potential punishment induced responses that were slower (*F*_(1,30)_=35.28, *p*<0.001) but *also* more accurate (*Chisq*_(1)_=26.73, *p*<0.001; Figure 2B). These results control for trial-to-trial differences in congruence, which, as expected, revealed faster (*F*_(1,31)_=115.28, *p*<0.001) and more accurate (*Chisq*_(1)_=4.13, *p*=0.042) responses for congruent stimuli compared to incongruent stimuli. Although there were no significant two-way interactions between incentives and congruency on performance, we observed a significant three-way interaction between reward, penalty, and congruence (*Chisq*_(1)_=6.24, *p*=0.013) specific to accuracy. Together, these data suggest that participants applied distinct strategies for engaging cognitive control across reward and punishment incentives.

**Table 2.** Mixed Model Results for Log-Transformed Reaction Time and Accuracy *: *p*<0.05, **: *p*<0.01, ***: *p*<0.001

| *Predictors* | **Log-transformed RT** |  |  | **Accuracy** |  |  |
| --- | --- | --- | --- | --- | --- | --- |
|  | *Estimates* | *S.E.* | *P-Value* | *Odds Ratios* | *S.E.* | *P-Value* |
| Age | 0.014 | 0.007 | 0.066 | 0.941 | 0.117 | 0.623 |
| Female - Male | -0.023 | 0.007 | **0.002\*\*** | 1.234 | 0.155 | 0.095 |
| High Penalty - Low Penalty | 0.014 | 0.002 | **<0.001\*\*\*** | 1.381 | 0.082 | **<0.001\*\*\*** |
| High Reward - Low Reward | -0.012 | 0.002 | **<0.001\*\*\*** | 1.028 | 0.039 | 0.464 |
| Trial Congruence (Cong-Incong) | -0.020 | 0.002 | **<0.001\*\*\*** | 1.105 | 0.050 | **0.028\*** |
| Reward X Penalty | -0.003 | 0.001 | **0.015\*** | 1.014 | 0.042 | 0.729 |
| Penalty X Congruence | 0.001 | 0.001 | 0.353 | 1.043 | 0.038 | 0.256 |
| Reward X Congruence | -0.001 | 0.001 | 0.432 | 1.044 | 0.039 | 0.249 |
| Reward X Penalty X Congruence | 0.000 | 0.001 | 0.543 | 1.097 | 0.041 | **0.012\*** |
| Number of Subjects | 32 |  |  | 32 |  |  |
| Observations | 27509 |  |  | 28785 |  |  |
| Marginal R<sup>2</sup> / Conditional R<sup>2</sup> | 0.056 / 0.255 |  |  | 0.055 / 0.150 |  |  |

### Reward Rate-Optimal Control Allocation: Normative Predictions

To generate predictions about performance on the Stroop task, we parameterized the tasks as a process of noisy evidence accumulating towards one of two boundaries (correct vs. error), using the *drift diffusion model* (DDM) [34,37]. We hypothesized that two of the DDM parameters that determine performance on a given trial are the rate of evidence accumulation (*drift rate, ν*) and the decision threshold (*a*). As the drift rate increases, the likelihood of a correct response increases (error rate decreases), and responses are *faster*. As the threshold increases, responses are also more likely to be correct but are *slower* (Figure 3A; [31]). As we describe below, a key prediction is that adjustments in these parameters may underlie divergent strategies for cognitive control allocation.

**Figure 3.**
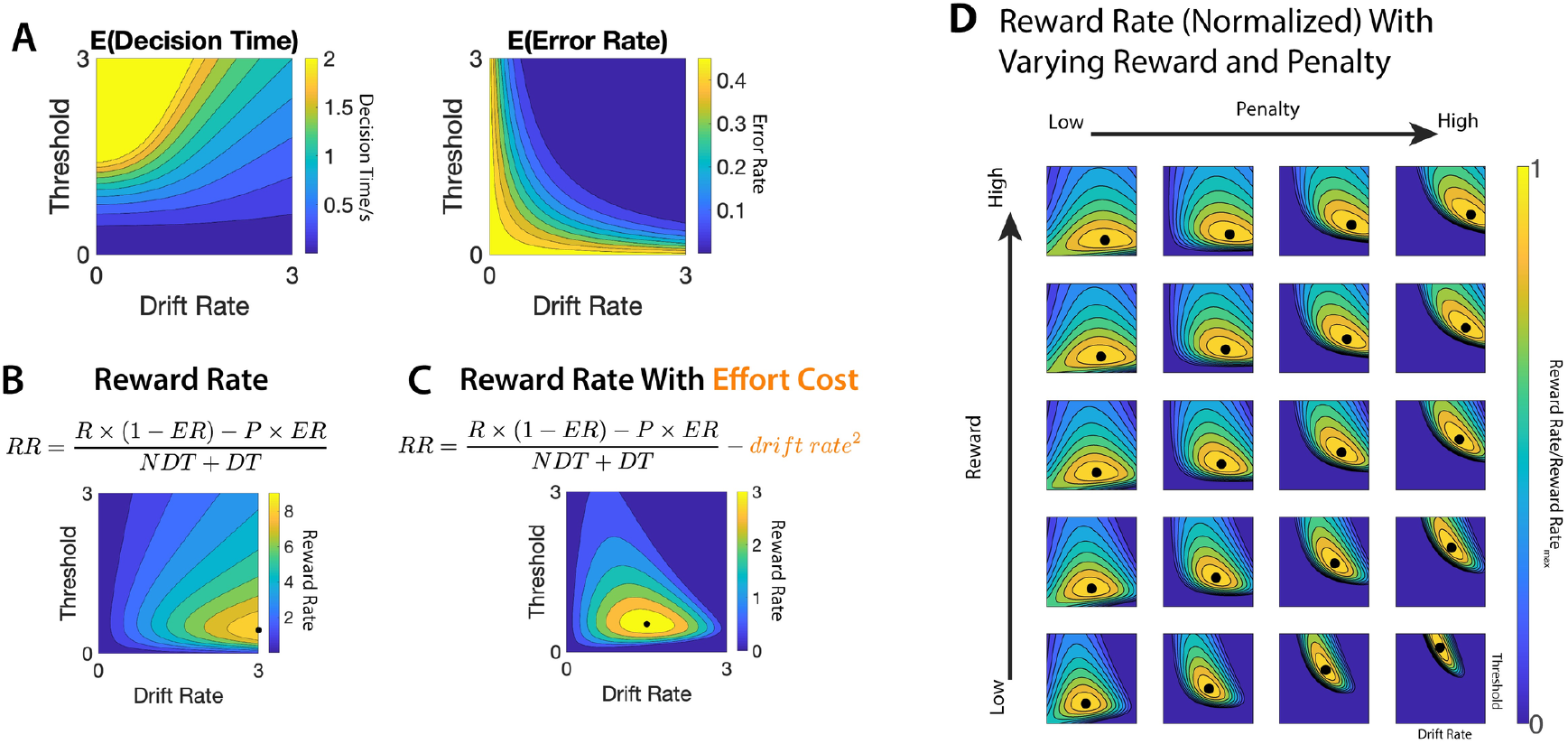
The influence of DDM parameter settings on estimates of reward rate. **A)** The expected error rate (*ER*) and decision time (*DT*) can be estimated as a function of drift rate and threshold. B-C) Reward rate is traditionally defined as a function of expected error rate, scaled by the value of correct vs. incorrect responses, and the overall response time (the combination of decision time and decision-unrelated processes [31]). The combination of drift rate and threshold settings that maximizes reward rate (black dots) differs depending on whether drift rate is assumed to incur an effort cost or not. Without a cost (B), it is always optimal to maximize drift rate. With a cost (C), drift rate and threshold must both fall within a more constrained set of parameter values. Parameters for (B-C): *R* = 5,*P* = 5,*NDT* = 0.4*s*. (D) As the reward for each correct response increases (from 8 to 20), the optimal joint configuration of drift rate and threshold (black dot) moves primarily in the direction of increasing drift rate. As the penalty for an incorrect response increases (from 5 to 625), this optimal configuration moves in the direction of increasing threshold.

Previous theoretical and empirical work has shown that participants can adjust parameters of this underlying decision process to maximize the rate at which they are rewarded over the course of an experiment [31,38]. This reward rate (*RR*) is determined by a combination of performance metrics (response time and error rate [*ER*], [31]) and the incentives for performance (i.e., outcomes for correct vs. incorrect responses):

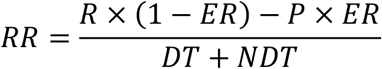

Here, the numerator (expected reward) is determined by the likelihood of a correct response (1 − *ER*), scaled by the reward for a correct response (*R*), relative to the likelihood of an error (*ER*), scaled by the associated punishment (*P*) [39]. The denominator (response time) is determined by the time it takes to accumulate evidence for a decision (decision time [*DT*]) as well as additional time to process stimuli and execute a motor response (non-decision time [*NDT*]).

To correctly respond to a Stroop trial (i.e., name stimulus color), participants need to recruit cognitive control to overcome the automatic tendency to read the word [40,41]. Building on past work [31,38,39], we can use the reward rate formulation above to identify how participants should normatively allocate control to maximize the reward rate (Figure 3B-C). To do so, we make three key assumptions. First, we assume that participants performing our task choose between adjusting two strategies for increasing their reward rate: (1) increasing attentional focus on the Stroop stimuli (resulting in increased drift rate toward the correct response), and (2) increasing their threshold to require more evidence accumulation before responding. Second, we assume that participants seek to identify the combination of these two DDM parameters that maximize reward rate. Third, we assume that increasing the drift rate incurs a nonlinear cost, which participants seek to minimize. The inclusion of this cost term is motivated by previous psychological and neuroscientific research [1] and by its sheer necessity for constraining the model from seeking implausibly high values of drift rate (i.e., as this cost approaches zero, the reward-rate-maximizing drift rate approaches infinity, as shown in Figure 3B). While a quadratic cost term was chosen a priori based on previous work [33,42], follow-up analyses (See Supplementary Results 1) indicated that the predictions made by this quadratic function are also more consistent with our data than those for a linear (i.e., absolute) function.

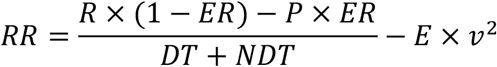

In this formula, *E*represents the weight of effort cost. Since the optimal drift rate and threshold are determined by the ratios *R/E* and *P*/*E*, the magnitude of effort costs is held constant (*E* = 1) for the reward rate optimization process, putting reward and punishment into units of effort cost. With this modified form of reward rate, the optimal drift rate is well-constrained (Figure 3C).

Using this formulation of reward rate (*RR*), we can generate predictions about the allocation of cognitive control (the combination of drift rate and threshold) that would be optimal under different reward and punishment conditions. To do so, we varied reward and punishment values and, for each pair, identified the pair of drift rate and threshold that would maximize reward rate. As reward increases, the model suggests that the optimal strategy is to increase the drift rate. As punishment increases, the optimal strategy is to increase the threshold (Figure 4A). These findings indicate that the weights for rewards and punishments jointly modulate the optimal strategy for allocating cognitive control and that these two types of incentives focus on distinct aspects of the strategy. Specifically, they predict that people will tend to increase drift rate the more they value receiving a reward for a correct response. In contrast, people will adjust their threshold depending on how much they value receiving a reward for a correct response (decrease threshold) and receiving a punishment for an incorrect response (increase threshold).

**Figure 4.**
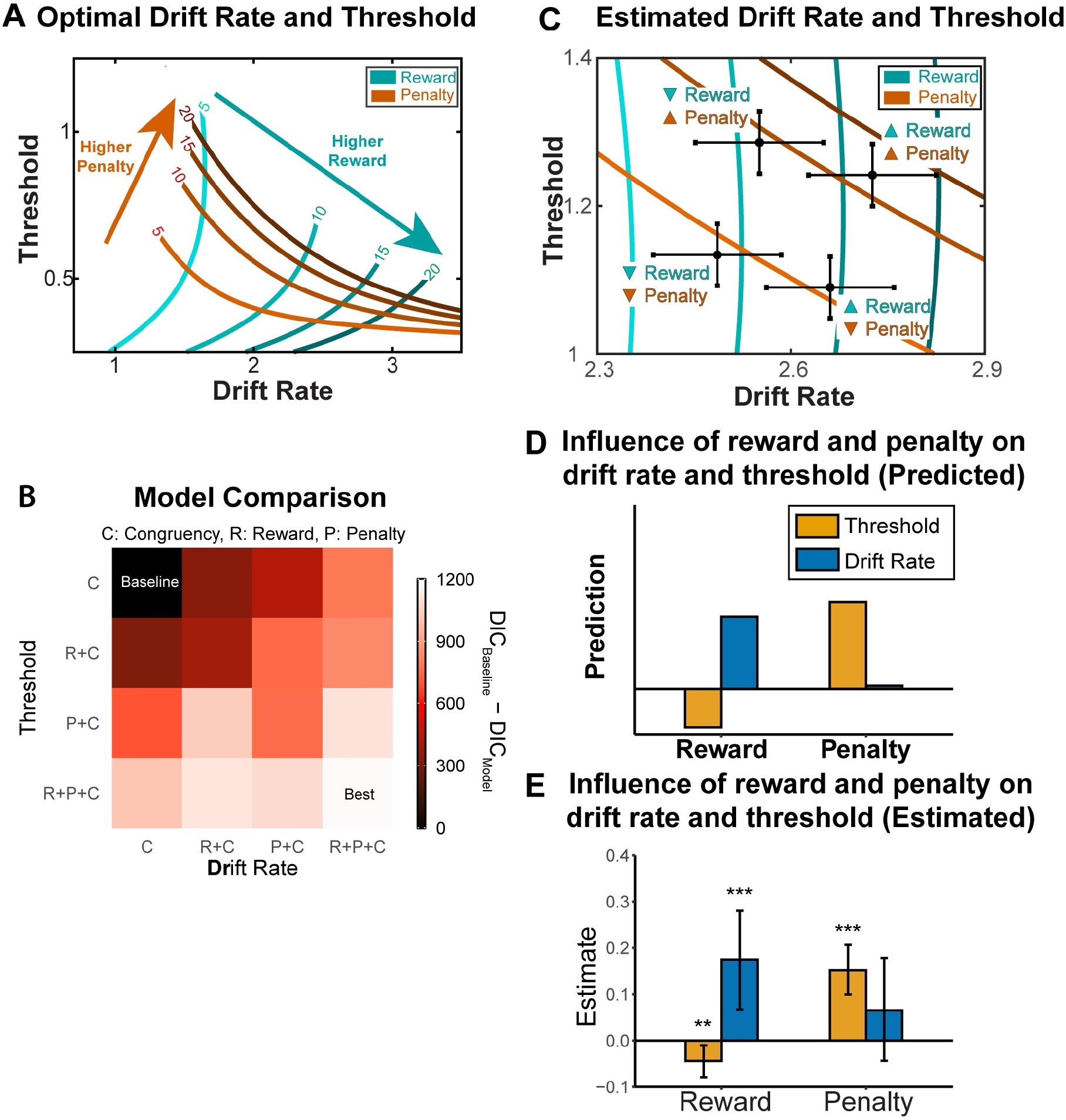
Normative and empirically observed estimates of incentive effects on DDM parameters. **A)** Combinations of drift rate and threshold that optimize (cost-discounted) reward rate, under different values of reward and penalty. **B)** We fit our behavioral data to different parameterizations of the DDM, with drift rate and/or threshold varying with reward, penalty, and/or congruence levels. The best-fitting model varied both DDM parameters with all three task variables. **C)** Estimated combination of drift rate and threshold for four conditions in the experiment. Error bars reflect s.d. **D-E)** Consistent with predictions based on reward-rate optimization (**D**, cf. panel **A**), we found that larger expected rewards led to increased drift rate, where as larger expected penalties led to increased threshold (**E**, cf. panel **C**). To a lesser extent, we found a decreased threshold with higher expected rewards. Error bars reflect 95% CI. *: p<0.05; ***: p<0.001. See also Figure A5 in S1 Supporting Information.

### Reward Rate-Optimal Control Allocation: Empirical Evidence

To test whether task performance was consistent with the predictions from our normative model, we fit behavioral performance on our task (reaction time and accuracy) with the Hierarchical Drift Diffusion Model (HDDM) package [43]. A systematic model comparison showed that the best-fitting parameterization of this model for our task allowed both drift rate and threshold to vary with trial-to-trial differences in congruency, reward level, and/or penalty level (Figure 4B; also see Supplementary Results 2). Critically, the parameter estimates from this model were consistent with predictions of our reward rate-optimal DDM (Figure 4C-E). Consistent with normative predictions, we found that reward and punishment exhibited dissociable influences on DDM parameters, such that larger rewards increased drift rate and decreased threshold, whereas larger punishment promoted a higher threshold. These findings control for the effect of congruency on DDM parameters (with incongruent trials being associated with lower drift rate and higher threshold). Taken together, our empirical findings are consistent with the prediction that participants are optimizing reward rate, accounting for potential rewards, potential punishments, and effort costs.

### Inferring Individual Differences in Sensitivity to Reward and Punishment

Our findings show that performance varies as a function of expected reward and punishment, and that these performance changes are consistent with a normative model according to which participants are maximizing reward and minimizing effort costs. However, both our model predictions and empirical findings also show that performance alone is insufficient to determine to what extent a participant was driven by a given incentive. For instance, faster performance could result from a participant being more sensitive to rewards, less sensitive to penalties, or both. The same is even true for estimates of individual model parameters within each of these conditions - our model predicts that a more reward-sensitive participant will lower their threshold than a less reward-sensitive participant, but that the same would be true for participants less vs. more sensitive to penalties. However, a key feature of our normative model is that it predicts how people will *jointly* configure control over drift rate and threshold based on their expected reward rate in a given condition, and predicts unique *combinations* of these DDM parameters under a given level of expected reward and penalty (Figure 4A). As a result, we can examine how participants move across this two-dimensional space as their rewards and penalties vary (Figure 5A), in order to make more robust inferences about the extent to which their performance was driven by each of these incentives. In other words, we can “reverse-engineer” how sensitive that participant had been to the rewards and penalties associated with performance on our task.

**Figure 5.**
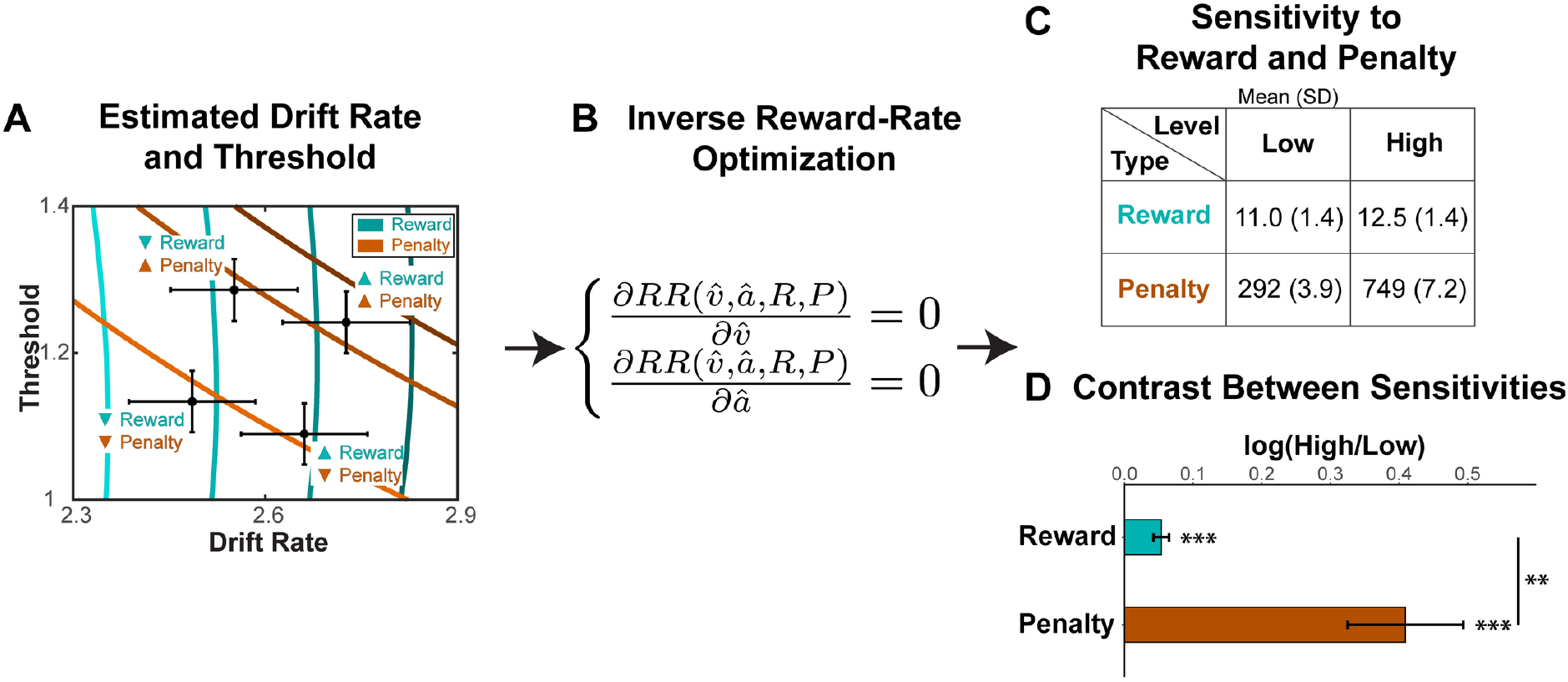
Inference of sensitivity to reward and penalty based on DDM estimates and reward rate optimization model. **A)** Estimated group-level reward-rate optimal combinations of drift rate and threshold for the four conditions in the experiment. Error bars reflect s.d. **B)** To infer the sensitivity to reward and penalty for a given individual, we invert this reward-rate optimization procedure, estimating the set of reward and penalty weights (*R* and *P*) that best accounts for that person’s pattern of behavior in a given condition. **C-D)** The resulting estimates of sensitivity to reward and penalty recapitulate our experimental manipulation, with higher sensitivity to reward in the high vs. low reward condition, and higher sensitivity to penalty for the high vs. low penalty condition. Panel **C** shows summary statistics across individual participants. Panel **D** shows a summary of individual-level contrasts between sensitivity to high vs. low reward and penalty. Error bars reflect s.e.m. **: p<0.01; ***: p<0.001. Parameter recovery validates subjective weight estimates (see Figure A7 in S1 Supporting Information).

To accomplish this, we used inverse reward-rate optimization to infer the individualized subjective weights of reward and punishment across the four task conditions based on participants’ estimated DDM parameters. For each task condition, we first estimated the drift rate (*ν*) and threshold (*a*) for each individual. We then calculated the partial derivatives of reward rate (*RR*) with respect to these condition-specific estimates of *ν* and *a*. By setting these derivatives to 0 (i.e., optimizing the reward-rate equation), we can calculate the sensitivity to reward and punishment (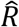 and 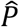) that make the estimated DDM parameters the optimal strategy (Figure 5C). This workflow can be summarized as follows:

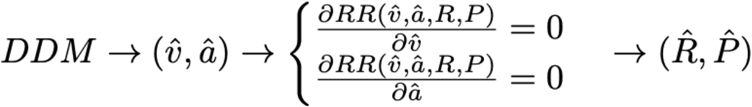

To validate this approach, we simulated DDM parameters under different combinations of reward and penalty sensitivities (*R* and *P*), and tested whether we could recover the ground-truth parameters based on simulated data. We were able to successfully recover both of these parameters (Part 5 in S1 Supporting Information; correlation between simulated and recovered values: r = 0.99 for *R*, and r = 0.93 for *P*), confirming that our estimation approach can be effective at inferring individual’s subjective valuation of reward and punishment when determining cognitive control adjustments.

A repeated-measures ANOVA on our estimates of *R* and *P* (log-transformed) revealed a main effect of incentive magnitude (*F*_(1,251)_=12.64, *p*=4.5e-4), with larger 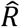 on high-reward intervals (*t*_(31)_=4.9, *p*=3.2e-5) and larger 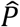 on high-punishment intervals (*t*_(31)_=4.72, *p*=4.8e-5). We also observed a main effect of valence, such that estimates of 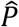 were higher than estimates of 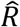 *F*_(1,251)_=603.70, *p*<2e-16). The ANOVA also revealed a significant interaction between valence and magnitude (*F*_(1,251)_=7.47, *p*=0.007; see Figure 5D), such that 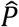 estimates differed more across punishment levels than 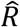 estimates differed across reward levels. These asymmetric effects of rewards and punishment on reward rate are consistent with research on loss aversion [44] and error aversion [45].

### Replication and extension of Study 1 findings in an independent sample

To verify the robustness of our observed dissociation between reward effects on drift rate and penalty effects on threshold, we recruited a separate group of participants (N=65) to perform our task. To further investigate whether these effects generalize beyond two levels of reward and penalty, we also included an intermediate level of reward and penalty between the two extremes previously tested. The magnitude of reward and punishment in each interval was therefore selected independently from three possible levels: 1 cent (Low), 5 cents (Medium) and 10 cents (High). The selected reward and punishment are then combined into a cue indicating these incentive levels.

This second study replicated the dissociable behavioral patterns observed in Study 1. Consistent with the previous study, we found that participants were faster (*F*_(2,64)_=13.91, *p*<0.001) but similarly accurate (*Chisq*_(2)_=2.23, *p*=0.317) with higher levels of reward, resulting in an overall higher number of correct responses per second as expected reward increased (*F*_(2,70)_=12.28, *p*<0.001; Figure 6A). Also consistent with Study 1, participants were slower (*F*_(2,63)_=8.49, *p*<0.001) but more accurate (*Chisq*_(2)_=15.21, *p*<0.001) with higher levels of punishment, resulting in fewer correct responses per second (*F*_(2,64)_=4.30, *p*=0.018; Figure 6B). Response rates under Medium levels of reward and penalty were intermediate to response rates under Low and High levels of those respective variables.

**Figure 6.**
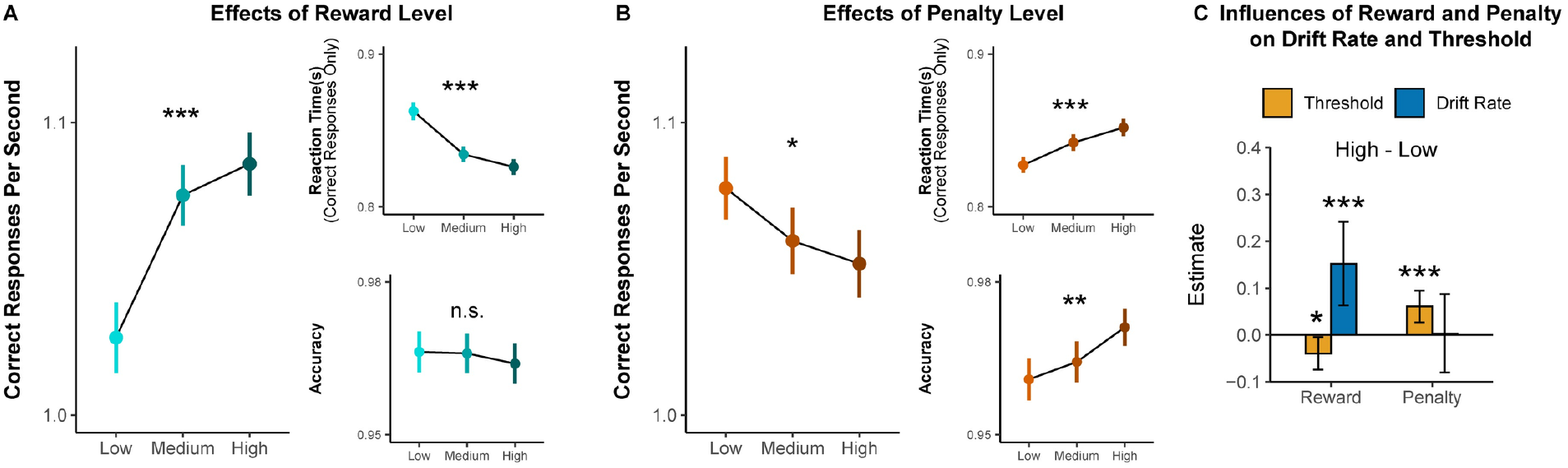
Effects of reward and punishment on overall task performance (A,B) and parameters of drift diffusion model (C) in Study 2. **A)** With increasing expected reward, participants completed more correct responses per second within a given interval (**Left**), which reflect faster responding on correct trials (**top right**) without any change in overall accuracy (**bottom right**). **B)** With increasing expected punishment, participants instead completed fewer trials per second over an interval, reflecting slower and more accurate responses. **C)** Drift rate increases with higher expected reward while threshold increases with higher expected punishment. Error bars reflect 95% CI. n.s.: p>0.05; *: p<0.05; **: p<0.01; ***: p<0.001.

When fitting Study 2 data with our best-fitting model from Study 1, we replicate the normatively predicted dissociation observed in that study. Reward exerted a significant positive influence on drift rate (*p*<0.001) and negative influence on threshold (*p*=0.013). Penalty exerted a significant positive influence on threshold (*p*=0.008) but not drift rate (*p*=0.47). These findings are consistent with the predictions from the reward rate optimization model.

## Discussion

We investigated divergent influences of reward versus punishment on cognitive control allocation, and the normative basis for these incentive-related control adjustments. Participants performed a self-paced cognitive control task that offered the promise of monetary rewards for correct responses and penalized monetary losses for errors. We found that higher potential rewards led to faster but equally accurate responding (resulting in increased monetary earnings), whereas higher potential punishment led to more accurate but slower responding (thus earning less reward but avoiding punishment). We showed that these dissociable patterns of incentive-related performance could be accounted for by two distinct strategies (adjustment of the strength of attention vs. response threshold), which are differentially optimal (i.e., reward rate maximizing) in response to these two types of incentives.

Our findings build on past research on reward rate maximization that has shown that people flexibly recruit cognitive control to maximize their subjective reward per unit time [30,31,35]. Our current experiments build on this research in several important ways. First, we apply this reward rate optimization model to performance in a self-paced variant of a cognitive control task. Second, we model and experimentally manipulate the incentive value for a correct versus incorrect response. Third, we incorporate the well-known cost of cognitive effort [1,46] into the reward rate optimization model (see below). Finally, we used our model to perform reverse inference on our data, identifying the subjective weights of incentives that gave rise to performance on a given trial.

We showed that adjustments of threshold and drift rate can vary as a function of task incentives, which then drive adaptive adjustments in cognitive control. Notably, achieving this result required us to build in the assumption that increases in drift rate incur a cost, an assumption that is grounded in past research on mental effort [1,33]. In the absence of this cost, our reward rate model predicts that individuals should maintain a maximal drift rate across incentive conditions, which is inconsistent with our findings. However, while we have ruled out the possibility that drift rate is costless, the precise form of its cost function remains an open question. Follow-up simulations show that our assumed quadratic cost function -- which was motivated by previous research into cognitive effort discounting [47,48] -- offers a smoother objective function than linear or exponential alternatives (Figure A3 in S1 Supporting Information), but all three of these cost functions make qualitatively similar predictions for our current task. We have also left open the question of whether and how a cost function applies to increases in response threshold. While there is reason to believe that threshold adjustments may incur analogous effort costs to attentional adjustments, in part given the control allocation mechanisms they share [2,32,34,49–51], threshold adjustments already carry an inherent cost in the form of a speed-accuracy tradeoff. It therefore wasn’t strictly necessary to incorporate an additional effort cost for threshold in the current simulations (Figure A4 in S1 Supporting Information), though it is possible such a cost would provide additional explanatory power under a different task design. Future work should investigate potential differences in these cost functions across these and other common control signals.

While our modified reward rate optimization model was able to accurately characterize how reward and punishment incentives influenced cognitive control allocation in our task, a critical next step will be to examine the degree to which these findings generalize to other tasks and incentive schemes, and to refine the model accordingly. For instance, in addition to testing the form that different control cost functions take, future work can clarify how people discount time when optimizing this reward function. Our model assumes that people discount time in a multiplicative fashion (i.e., as the denominator for reward), which is a standard assumption in models of reward rate optimization [31,38]. However, we cannot rule out an alternative possibility that they are instead discounting time additively, as is assumed by models that treat time as an opportunity cost of effort [35,52], because these models are likely to make similar predictions with respect to drift and threshold optimization in our current study. Identifying and testing tasks that differentiate between these predictions holds value for bridging these two lines of research in the service of better understanding effort allocation.

Another open question is whether people weigh the incentives for a correct response differently depending on whether these incentives are positive or negative. In our study, correct responses were only associated with potential rewards (positive reinforcement), but a key prediction of our model is that people should adjust their control configuration similarly (i.e., increase drift rate, lower threshold) when correct responses instead avoid a negative outcome (negative reinforcement), though perhaps to different degrees. Our approach thus offers promise for disentangling the roles of incentive valence (positive vs. negative) and incentive type (reinforcement vs. punishment) in motivated control [53].

More generally, it will be important to test whether similar drift and threshold adjustments occur across other cognitive control tasks that carry a similar structure to this one, and to extend our optimization approach to tasks that require different forms of multivariate control configuration, such as distributing attention across multiple stimuli or features [54,55]. Broadening the applications of this approach to a wider array of control signals will also provide a critical step towards understanding how people distribute their cognitive effort across a multitude of tasks in real-world settings. Along these lines, a simplifying assumption of our current approach was that people assume reward rate is constant within a given task environment. While this assumption was reasonable given the parameters of our task (i.e., where incentives were explicitly cued and pseudorandomized), a crucial next step will be to examine how people dynamically reconfigure control as they learn from feedback that the expected rewards and penalties in their environment are changing. Research has shown that people dynamically adjust their response threshold in both decision-making tasks [56] and cognitive control tasks [30,57] as they learn to expect greater rewards. It remains to be tested how these cognitive control adjustments are distributed across both threshold and drift rate with changes in both reward and punishment, as well as with individual-specific [58,59] and context-specific [60] differences in learning from these positive and negative outcomes.

Interestingly, research into how people learn differentially from positive versus negative outcomes is that these learned values also differentially influence a person’s confidence on a given task, with negative feedback resulting in lower confidence in one’s performance on both perceptual and value-based choice tasks [61,62] Given the connections that have been separately drawn between confidence and adjustments of response threshold [63,64], these findings converge with our own observations of increasing threshold in the face of higher expected punishment. Thus, an important direction for future work will be to examine how metacognitive experiences associated with our task vary with experienced incentives and potentially serve to moderate subsequent control adjustments.

Finally, our combined theoretical and empirical approach enabled us to quantify individual differences in how participants subjectively valued expected rewards and punishments based solely on their task performance. We found that people weighed punishments more heavily than rewards, despite the equivalent currency (i.e., amounts of monetary gain vs. loss). This finding is consistent with past work on loss aversion [44] and motivation to avoid failure [45,65], and more generally, with the findings that distinct neural circuits are specialized for processing appetitive versus aversive outcomes [66,67]. While our approach to estimating these individual differences is exploratory and requires further validation across different tasks and incentive schemes (such as those noted above), we believe that it holds promise for understanding how people vary in their motivation to succeed and/or avoid failure in daily life [21,68–72]. Not only can this method help to infer these sensitivity parameters for a given individual implicitly (i.e., based on task performance rather than self-report), it can also provide valuable insight into the cognitive and computational mechanisms that underpin adaptive control adjustments, and when and how they become maladaptive (e.g., for individuals with anxiety, depression, or schizophrenia) [73–78].

## Materials and Methods

### Participants

#### Study 1

We collected 36 participants online through Amazon’s Mechanical Turk. We limited the sample to participants located within the United States, but did not put any other restrictions on demographics (e.g., race). Participants gave informed written consent and received cash ($3 to $6, depending on their performance and task contingencies) for participation. The study was approved by Brown University’s Institutional Review Board.

4 participants were excluded for either not understanding the task properly (based on their responses to quiz questions after the instructions) or having mean accuracy below 60% and mean reaction times outside of 3 standard deviations of the mean reaction time of all the participants. The remaining 32 participants (Gender: 31% Female; Age: 35±10 years) were included in all of our analyses.

#### Study 2

We collected 71 participants online through Amazon’s Mechanical Turk. Participants gave informed written consent and received cash ($3 to $6, depending on their performance and task contingencies) for participation. The study was approved by Brown University’s Institutional Review Board.

6 participants were excluded for either not understanding the task properly (based on their responses to quiz questions after the instructions) or having mean accuracy below 60% and mean reaction times outside of 3 standard deviations of the mean reaction time of all the participants. The remaining 65 participants (Gender: 45% Female; Age: 38±9 years) were included in all of our analyses.

### Incentivized Cognitive Control Task

#### Study 1

We designed a new task to investigate cognitive control allocation in a self-paced environment (Figure 1). During this task, participants are given fixed time intervals (e.g., 10 seconds) to perform a cognitively demanding task (Stroop task), in which they have to name the ink color of a color word. There were four possible ink colors (red, yellow, green and blue) across four possible color words (‘RED’, ‘YELLOW’, ‘GREEN’, ‘BLUE’). Participants were instructed to press the key corresponding to the ink color of each stimulus. The ink color could be congruent (e.g., BLUE) or incongruent (e.g., BLUE) with the meaning of the word. Responding to incongruent stimuli has been shown to require an override of their more automatic tendency to respond based on the word meaning. The overall ratio of congruent versus incongruent trials was 1:1. Participants could perform as many Stroop trials as they wanted and were able during each interval, with a new trial appearing immediately after each response. Due to this self-paced design, the proportion of congruent trials could vary slightly across intervals. To discourage participants from developing a trial-counting strategy (e.g., aiming to complete 10 responses per interval), the duration of intervals varied across the session (i.e., ranging from 8 to 12 seconds).

Participants were instructed that they would be rewarded for correct responses and penalized for incorrect responses. At the start of each interval, a visual cue indicated the level of reward and punishment associated with their responses in the subsequent interval. We varied reward for correct responses (+1 cent or +10 cents) and punishment for incorrect responses (-1 cent or -10 cents) within each subject, which leads to four distinct conditions (Figure 1). Each participant performed 20 intervals per condition. During the interval, participants could complete as many Stroop trials as they would like. Below each Stroop stimulus, a tracker indicated the cumulative amount of monetary reward within that interval. After each interval, participants were informed how much they earned. To ensure that each interval was evaluated independently, participants were informed (veridically) that 8 out of the 80 intervals in the main task were randomly selected and the total money earned in these selected intervals would be part of their final payment. The experiment was implemented within the PsiTurk framework [79].

Before the main task, participants performed several practice sessions. First, they practiced the mapping between keyboard keys and colors (80 trials). Then they completed practice for the Stroop task (60 trials). Participants then practiced the Stroop task in the self-paced setting (4 intervals). In a final practice block, participants were introduced to the visual cues and practiced the self-paced intervals with these visual cues (12 intervals).

#### Study 2

The task in Study 2 has a similar structure compared to Study 1. The major difference between tasks was that the magnitude of reward and penalty was selected from three possible levels (1 cent, 5 cents and 10 cents) instead of binary levels in Study 1, such that there exist 9 distinct conditions in the experiment (3 levels of reward by 3 levels of punishment, Figure 6). Same with Study 1, the condition was cued prior to the start of each interval.

### Analyses

#### Study 1

With this paradigm, we can analyze performance at the level of a given interval and at the level of responses to individual Stroop stimuli within that interval. We analyzed participants’ interval-level performance by fitting a linear mixed model (lme4 package in R; [80] to estimate the correct responses per second as a function of contrast-coded reward and punishment levels (High Reward = 1, Low Reward = -1, High Punishment =1, Low Punishment = -1) as well as their interaction. The models controlled for age, gender, and proportion of congruent stimuli, and using models with maximally specified random effects [81].

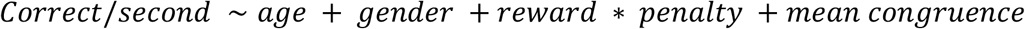

To understand how the incentive effects on overall performance are composed of the influences on speed and accuracy, we separately fit linear mixed models to trial-wise reaction time (correct responses only) and accuracy, controlling for the stimuli congruency. We performed analysis of variance on the fitted mixed models to test the overall effects of reward and punishment.

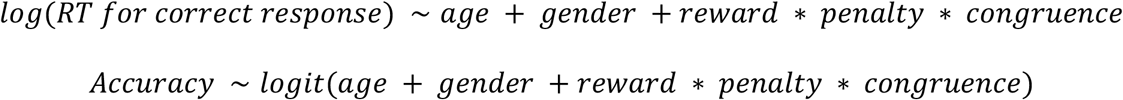

We parameterized participants’ responses in the task as a process of noisy evidence accumulating towards one of two boundaries (correct vs. error) using the Drift Diffusion Model (DDM). The DDM is a mechanistic model of decision-making that decomposes choices into a set of constituent processes (e.g., evidence accumulation and response thresholding), allowing precise measurement of how different components of the choice process (e.g., RT and accuracy) are simultaneously optimized [37]. We performed hierarchical fitting of DDM parameters using the HDDM package [43]. In the DDM model, the drift rate and threshold depend on trial type (congruent or incongruent), reward level and/or penalty level. The selection of predictors for drift rate and threshold is based on the model comparison using DIC. We fixed the starting point at the mid-point between the two boundaries as there was no prior bias toward a specific response in the task. The non-decision time was fitted as a free parameter.

We characterized the optimal allocation of cognitive control as the maximization of the reward rate [31] with modification for effort cost. Based on qualitative comparisons between predictions of different cost functions (Figures A3-A4 in S1 Supporting Information), we chose to express these cost functions as a quadratic function of drift rate and to assume no cost on increases in threshold, but note that alternate formats of each of these cost functions yield qualitatively similar predictions for all of our key findings (see Part 2 in S1 Supporting Information). With the effort-discounted reward rate, we make predictions about the influences of incentives on control allocation by numerically identifying the optimal drift rate and threshold under varying reward and punishment. To validate our normative prediction, we fit accuracies and RTs across the different task conditions with a DDM [43], which allowed us to derive estimates of how a participant’s drift rate and threshold varied across different levels of reward and punishment. We performed model comparison based on deviance information criterion (DIC; lower is better) to identify the best model for the behavioral data. Based on the assumption that participants’ cognitive control allocation optimizes the reward rate, we inferred participants’ subjective weights of reward and punishment from the estimated drift rate and threshold.

#### Study 2

We performed linear mixed model analysis on the participants’ interval-level performance with reward and punishment levels coded with sliding-difference contrast so that the two contrasts represent the difference between two consecutive reward or punishment levels (Medium - Low, High - Medium). We separately fit linear mixed models to trial-wise reaction time (correct responses only) and accuracy, controlling for the stimuli congruency.

We fit participants’ responses with the DDM using three-level polynomial contrast coding to obtain the linear and nonlinear patterns of incentive effects on DDM parameters. The coefficients in these contrasts were then transformed back to the DDM parameters under each condition.

All human data are available on OSF at link https://osf.io/24ud5/.

All code written in support of this publication is publicly available at https://github.com/Jasonleng/RewardPenaltyPaper.

## Acknowledgments

This work was funded by the Training Program for Interactionist Cognitive Neuroscience T32-MH115895 (X.L.), Training Program for Computational Psychiatry T32-MH126388 (D.Y.), an Innovation Award (A.S.) and a Daniel Cooper Graduate Student Fellowship (H.R.) from Brown’s Carney Institute for Brain Science; and by grants from the National Institute of General Medical Sciences (P20GM103645) and the National Science Foundation (CAREER Award 2046111) to A.S.

## Competing Interests

The authors have no competing interests to declare.

## Supporting Information Legend

**S1 Supporting Information. Supporting Information**.

## Notes

### Competing Interest Statement

The authors have declared no competing interest.

### Summary of Updates

updated discussion updated supporting information

