## Supplementary material for "Dissociable influences of reward and punishment on adaptive cognitive control": S1 Supporting Information

1. **Interaction between incentives and trial congruence**

In Study 1, We found that the influence of reward and penalty on behavioral performance was similar between congruent and incongruent trials (Figure A1A-B; see also Table 2). Similarly, additional mixed model analysis on data from Study 2 did not show significant interaction between either reward and trial congruence (reaction time: *F*_(2,246)_=1.32, *p*=0.27; accuracy: *Chisq*_(2)_=5.83, *p*=0.054; Figure A1C) or penalty and trial congruence (reaction Time: *F*_(2,63)_=1.54, *p*=0.22; accuracy: *Chisq*_(2)_=5.03, *p*=0.081; Figure A1D). We performed additional model comparison regarding the necessity of adding interaction between incentives and trial type into the model (Figure A2A). The best one includes an interaction between penalty and trial type for both drift rate and threshold. When looking at the interaction between penalty and trial type, we found significant effects on drift rate in both studies. In Study 1, we found that higher penalty increases the drift rate on incongruent trials but marginally decreased drift rate on congruent trials (Figure A2B). The interaction between penalty and congruence for drift rate in Study 2 follows the same direction in Study 1 (Figure A2C-D).

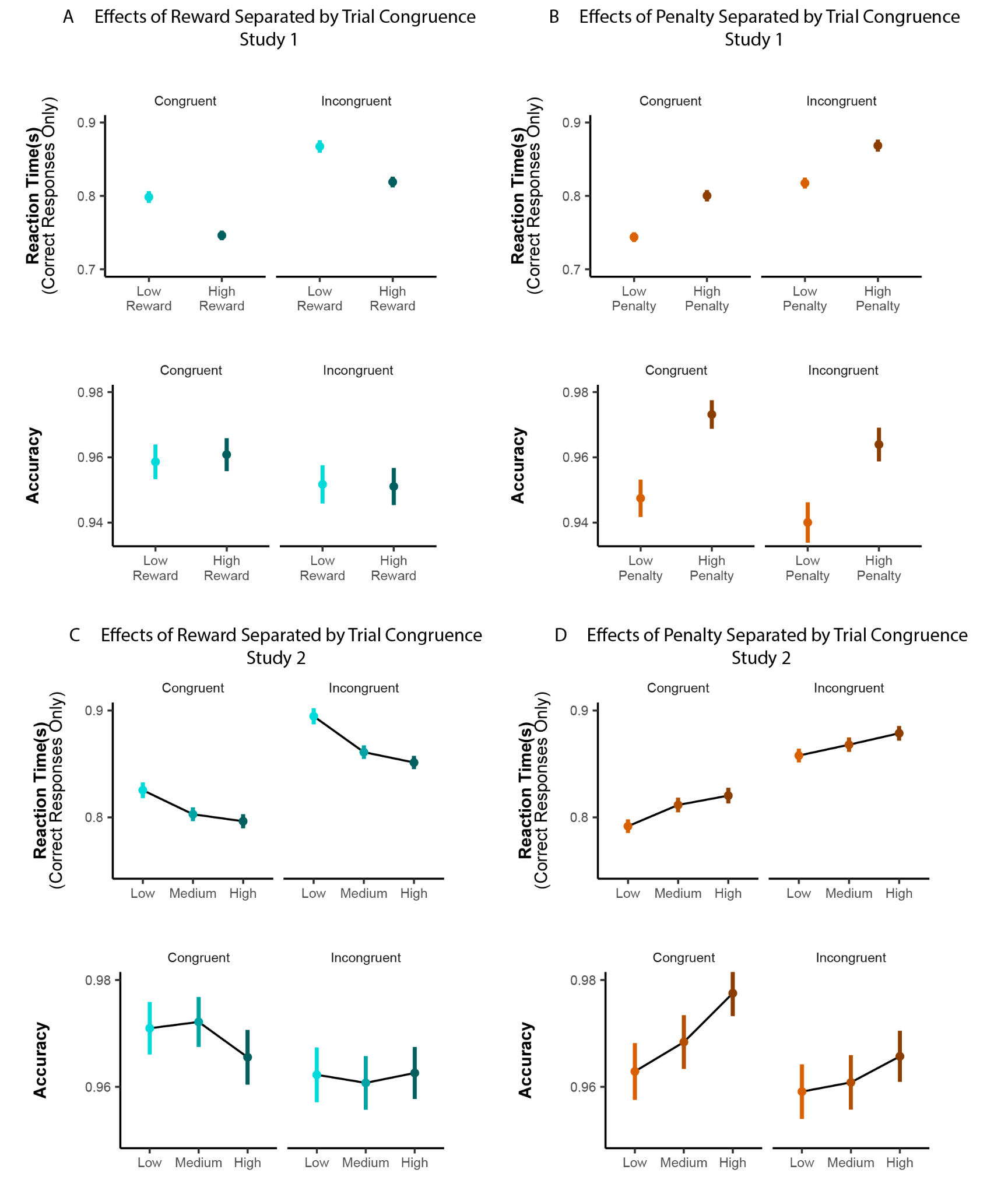

**Figure A1. Effects of reward and penalty in congruent and incongruent trials (A-B: Study 1; C-D: Study 2).** The effects of reward and penalty on reaction time and accuracy are consistent across congruent and incongruent trials.

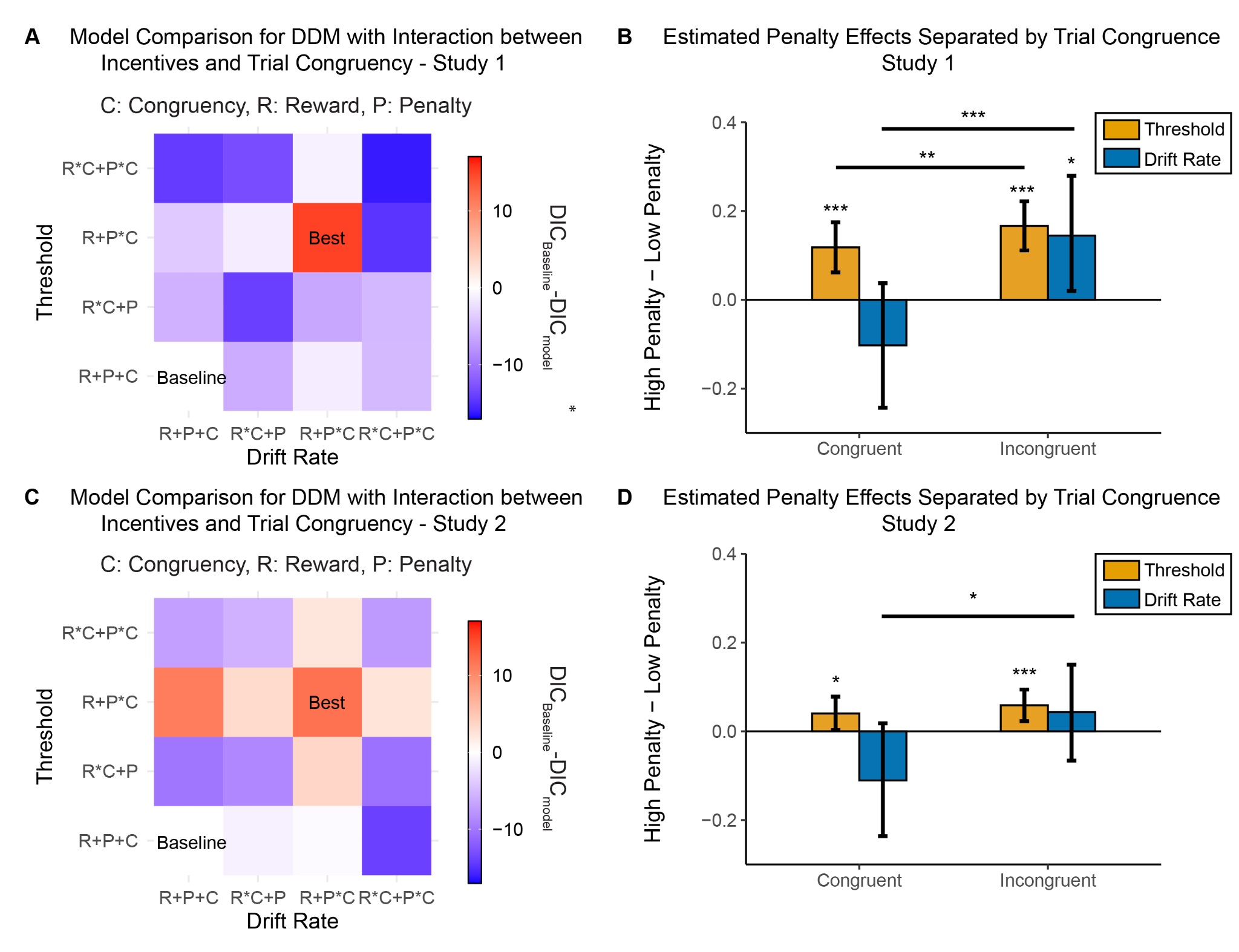

**Figure A2. Drift diffusion model with interaction between incentive level and trial congruence.** Model comparison based on DIC shows that the best model for Study 1 includes interaction between penalty and trial type for both drift rate and threshold **(A)**. The estimates of penalty effects separated by trial type in Study 1 is shown in **(B)**. Model comparison confirmed that the best model for Study 2 takes the same structure **(C)**. The estimates of penalty effects separated by trial type in Study 2 is shown in **(D).** Error bars reflect 95% CI. *: p<0.05; ***: p<0.001.

2. **Alternate effort cost functions for drift rate and threshold**

We considered linear, quadratic and exponential functions for effort cost of drift rate (denoted as $v$):

$$Linear, drift rate only: Cost \sim v$$

$$Quadratic, drift rate only: Cost \sim v^{2}$$

$$Exponential, drift rate only: Cost \sim e^{v}$$

The reward rate models with linear and exponential cost functions predict a discontinuity in the relationship between reward and drift rate (Figure A3A). This prediction is due to the bimodal profile of the reward rate function with these two alternatives, and the presence of a transition between modes when reward increases (Figure A3B). Based on these considerations and previous support for quadratic effort costs [1–5], we decided to use the quadratic function for our key model simulations. Critically, none of our predicted incentive effects differ substantially under either of these alternate effort cost functions.

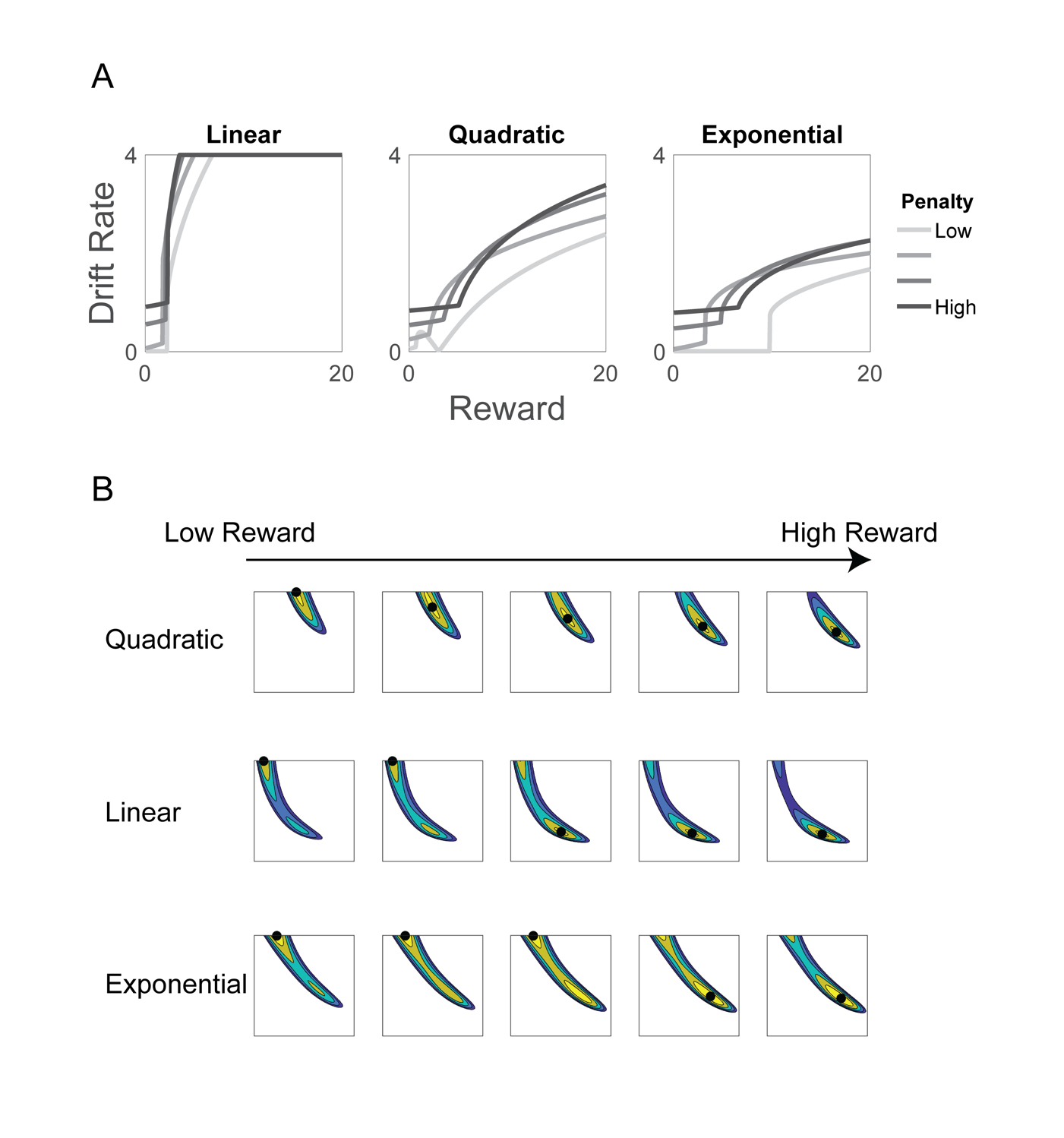

**Figure A3. Predicted relationship between reward and optimal drift rate with different format of effort cost.** Both linear and exponential formats of effort cost predict step function in the relationship between reward and optimal drift rate **(A)**. This is due to the existence of bimodality in the objective function **(B)**. A small increase of reward is associated with a shift of the optimal configuration of drift rate and threshold between the two modes.

We also considered the possibility that threshold (denoted as $a$) carried an effort cost rather than omitting such a cost (as in our prediction in the main text):

$$Quadratic, drift rate and threshold: Cost \sim v^{2}+a^{2}$$

We simulated how drift rate and threshold would jointly vary as a function of reward and penalty, both with and without an assumed threshold cost (in both cases assuming that drift rate carres a quadratic cost). The predictions of these two models were qualitatively very similar (Figure A4A-B), largely because threshold already carries a time cost for slower responding. The only distinguishing feature between these was the prediction of increased drift rate with higher penalties when assuming a cost for threshold (because these costs now trade off against the cost on drift rate), a prediction that our studies are unable to conclusively support or refute (Figure A4C). In the absence of clear evidence for this distinct prediction, we opted to omit a threshold cost for our main model simulations for the sake of parsimony but, as with alternate drift rate cost functions, none of our key conclusions would change under this alternative.

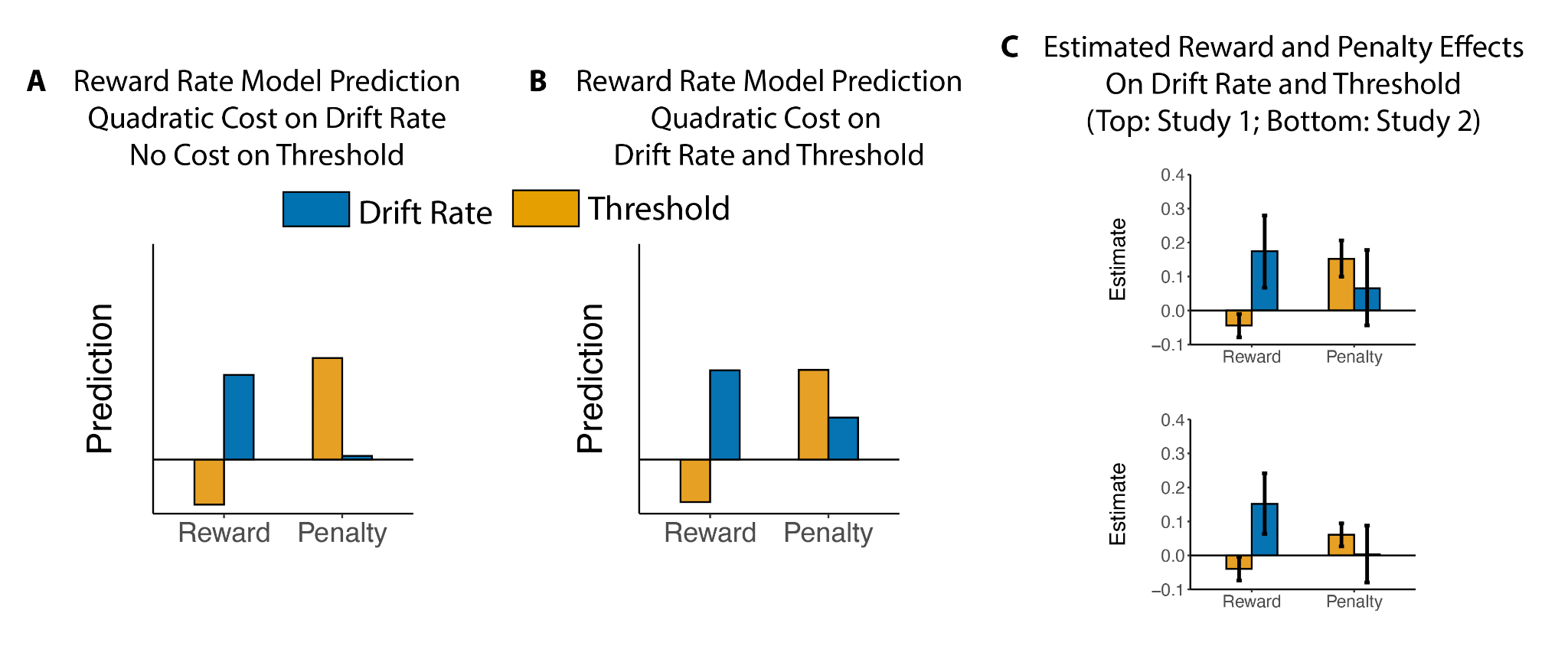

**Figure A4. Predicted incentive effects with and without effort cost of threshold.** The predicted reward and penalty effects on drift rate and threshold from reward rate model without cost on threshold **(A)** and with cost on threshold **(B)** are qualitatively consistent. The estimated reward and penalty effects in the current data **(C; adapted from Figure 4 and Figure 6)** do not provide strong evidence regarding whether the cost of threshold should be included in the model. Error bars reflect 95% CI.

3. **Model comparisons to DDM’s with task-modulated non-decision time**

In addition to the models estimated in Figure 4, we also considered models that estimated parameters for reward and penalty influences on non-decision time (NDT). Based on the best model shown in Figure 4B, we performed additional model comparison (Figure A5A). In the best model, we found a significant effect of penalty on NDT (Figure A5C). The best model with NDT maintains the same patterns of incentive effects on drift rate and threshold estimated in Figure 4, so that the findings in the paper are not due to incentive effects on stimuli encoding and/or motor responses. Based on these results, we did not include models with NDT in the paper.

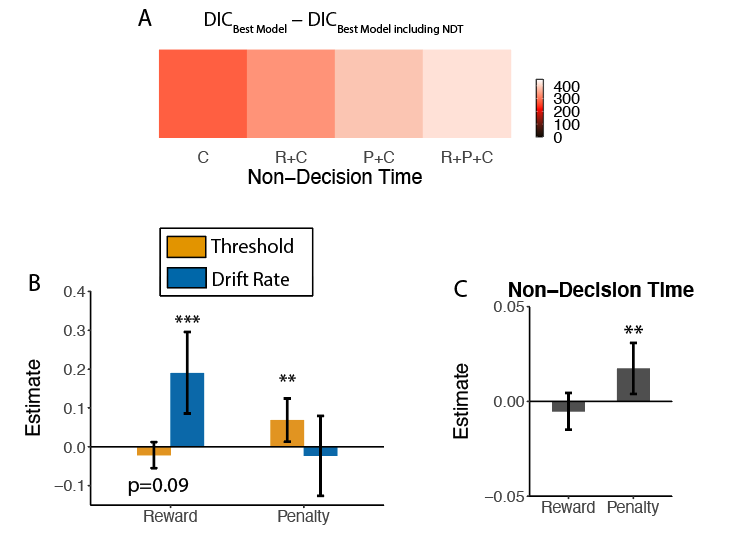

**Figure A5. Model comparison between best model in Figure 4B and models including non-decision time (Study 1).** **A)** The best model includes all predictors (reward, penalty and stimuli congruency) for non-decision time. **B)** Estimated effect of reward and penalty on threshold, drift rate and non-decision time for the selected model from Figure A2A. The patterns in threshold and drift rate are consistent with the best model in Figure 4B. **C)** There is a significant effect of penalty (but not of reward) on non-decision time. Error bars reflect 95% CI. **: p<0.01; ***: p<0.001.

4. **Posterior predictive check for Drift Diffusion Model (DDM)**

We performed a posterior predictive check for the fitted Drift Diffusion Model (DDM). We generated 500 independent samples from the posterior distribution of fitted parameters and then simulated the reaction time distribution for each posterior sample. The predicted reaction time distribution matches with the actual reaction time distribution for each condition (Figure A6).

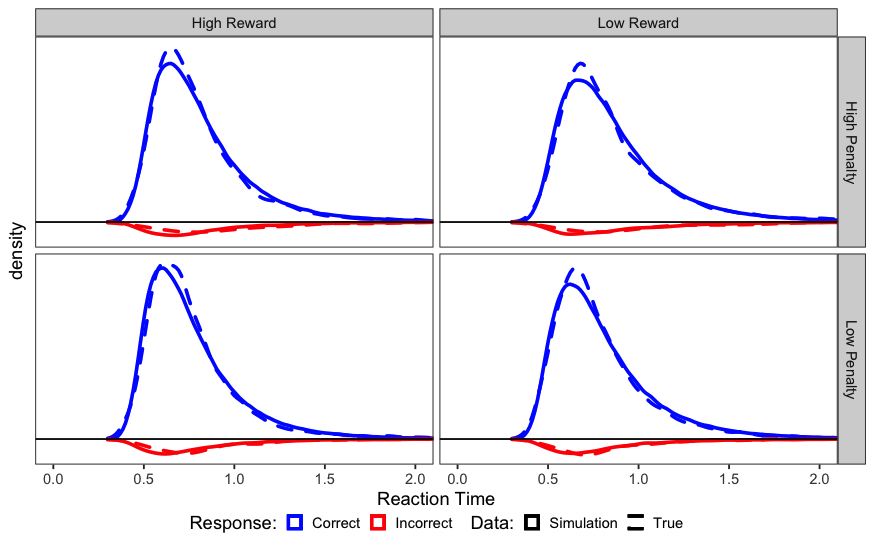

**Figure A6. Posterior predictive check for DDM (Study 1).** For each condition, the distribution of simulated reaction time from posterior predictive check (solid line) matches with the actual reaction time distribution (dashed line) for both correct and incorrect responses.

5. **Parameter recovery for subjective weights of reward and penalty**

We performed simulation-based parameter recovery for subjective weights of reward and penalty. To cover the range of inferred reward and penalty values, we sampled uniformly in [4,10] and [10,70] independently for reward and penalty to create 100 pairs of reward and penalty values. The non-decision time for each pair of reward and penalty is sampled uniformly in [0.35,0.45]. For each pair of reward and penalty values, we find the optimal drift rate and threshold that optimize the reward rate. We then generate 2000 trials for each pair of reward and penalty. By fitting the distribution of reaction time and accuracy with drift diffusion model, we estimate the drift rate, threshold and the non-decision time. Using these estimated parameters, we performed the inverse reward-rate optimization to infer the reward and penalty values (Figure A7). Both reward and penalty can be recovered from the inverse reward-rate optimization process.

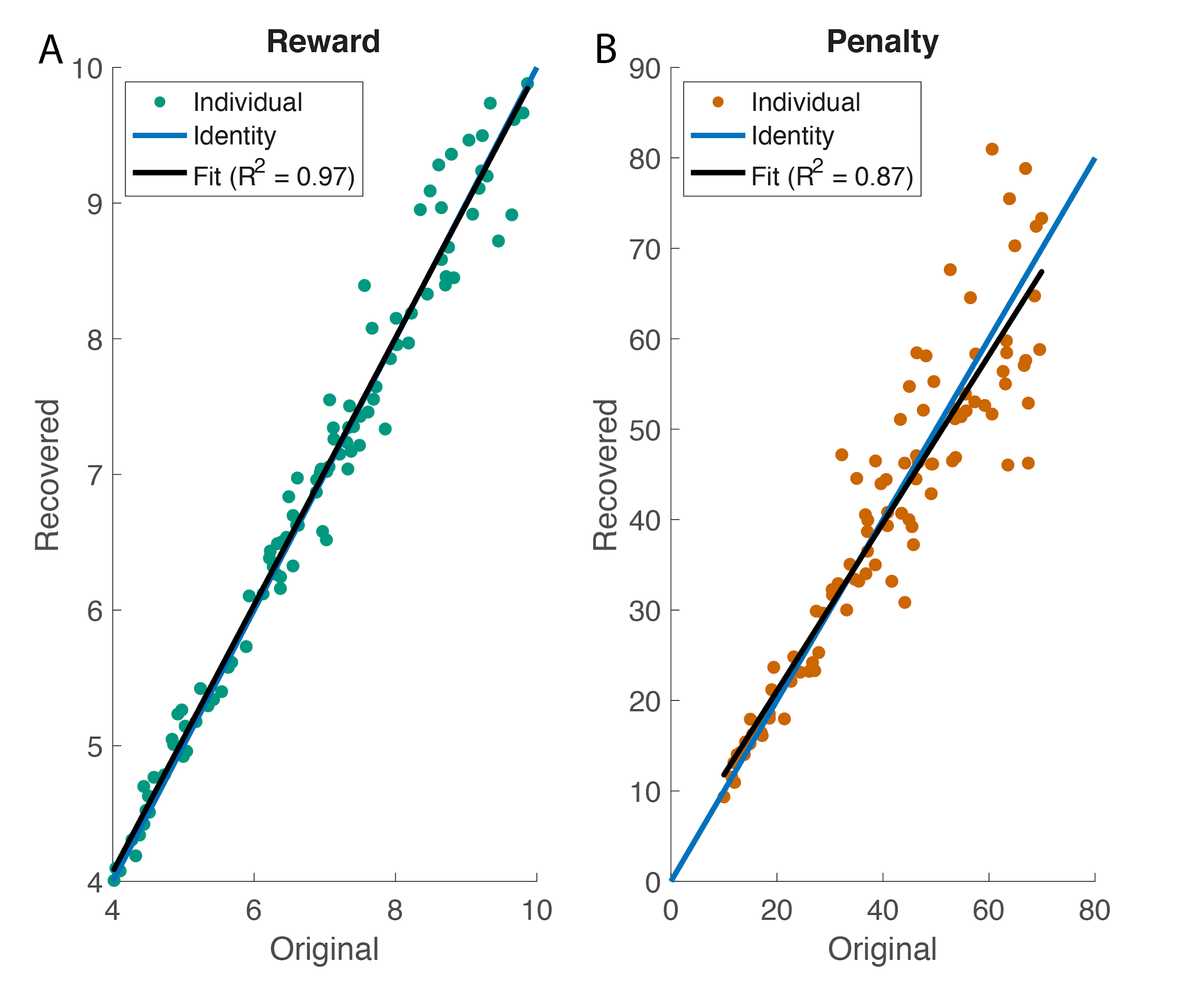

**Figure A7. Parameter recovery for the subjective weights (Study 1).** **A)** recovered reward versus original reward. **B)** recovered penalty versus original penalty. Both reward and penalty can be recovered from the inverse reward-rate optimization process.

6. **Mixed model Results for behavioral data from Study 2**

Table A1 and A2 show the details of the fitted mixed models for response rate, reaction time and accuracy.

**Table A1. Mixed Model Results for Correct Responses per Second**

**(Study 2)**

*: *p*<0.05, **: *p*<0.01, ***: *p*<0.001

*Italic*: does not reach threshold of significance but is trending

|  | **Correct Responses Per Second** | | |
| --- | --- | --- | --- |
| *Predictors* | *Estimates* | *S.E.* | *P-Value* |
| Age | -0.017 | 0.031 | 0.586 |
| Female - Male | -0.034 | 0.030 | 0.256 |
| Medium Reward - Low Reward | 0.051 | 0.010 | **<0.001***** |
| High Reward - Medium Reward | 0.009 | 0.008 | 0.254 |
| Medium Penalty - Low Penalty | -0.020 | 0.008 | **0.013*** |
| High Penalty - Medium Penalty | -0.008 | 0.008 | 0.325 |
| Average Congruence | 0.002 | 0.003 | 0.502 |
| Number of Subjects | 65 | | |
| Observations | 4490 | | |
| Marginal R^2^ / Conditional R^2^ | 0.028 / 0.675 | | |

**Table A2. Mixed Model Results for Log-Transformed Reaction Time and Accuracy**

**(Study 2)**

*: *p*<0.05, **: *p*<0.01, ***: *p*<0.001

*Italic*: does not reach threshold of significance but is trending

|  | **Log-transformed RT** | | | **Accuracy** | | |
| --- | --- | --- | --- | --- | --- | --- |
| *Predictors* | *Estimates* | *S.E.* | *P-Value* | *Odds Ratios* | *S.E.* | *P-Value* |
| Age | 0.010 | 0.008 | 0.197 | 1.110 | 0.085 | 0.173 |
| Female - Male | 0.008 | 0.007 | 0.262 | 1.081 | 0.080 | 0.297 |
| Medium Reward - Low Reward | -0.012 | 0.002 | **<0.001***** | 0.862 | 0.067 | *0.055* |
| High Reward - Medium Reward | -0.003 | 0.002 | *0.073* | 0.987 | 0.071 | 0.857 |
| Medium Penalty - Low Penalty | 0.007 | 0.002 | **0.001**** | 1.083 | 0.078 | 0.268 |
| High Penalty - Medium Penalty | 0.004 | 0.002 | *0.073* | 1.296 | 0.119 | **0.002**** |
| Trial Congruence (Cong-Incong) | -0.016 | 0.001 | **<0.001***** | 1.089 | 0.045 | **0.039*** |
| Number of Subjects | 65 | | | 65 | | |
| Observations | 35861 | | | 36955 | | |
| Marginal R^2^ / Conditional R^2^ | 0.044 / 0.441 | | | 0.024 / 0.139 | | |

7. **Drift diffusion model results for Study 2**

We fit the behavioral performance (reaction time and accuracy) in Study 2 with the HDDM with the selected model from Study 1. We found that drift rate increased with larger rewards (low to medium and medium to high), and threshold increased with larger punishment (medium to high). These results are consistent with normative prediction and previous findings in Study 1 that reward and punishment exhibited dissociable influences on drift rate and threshold.

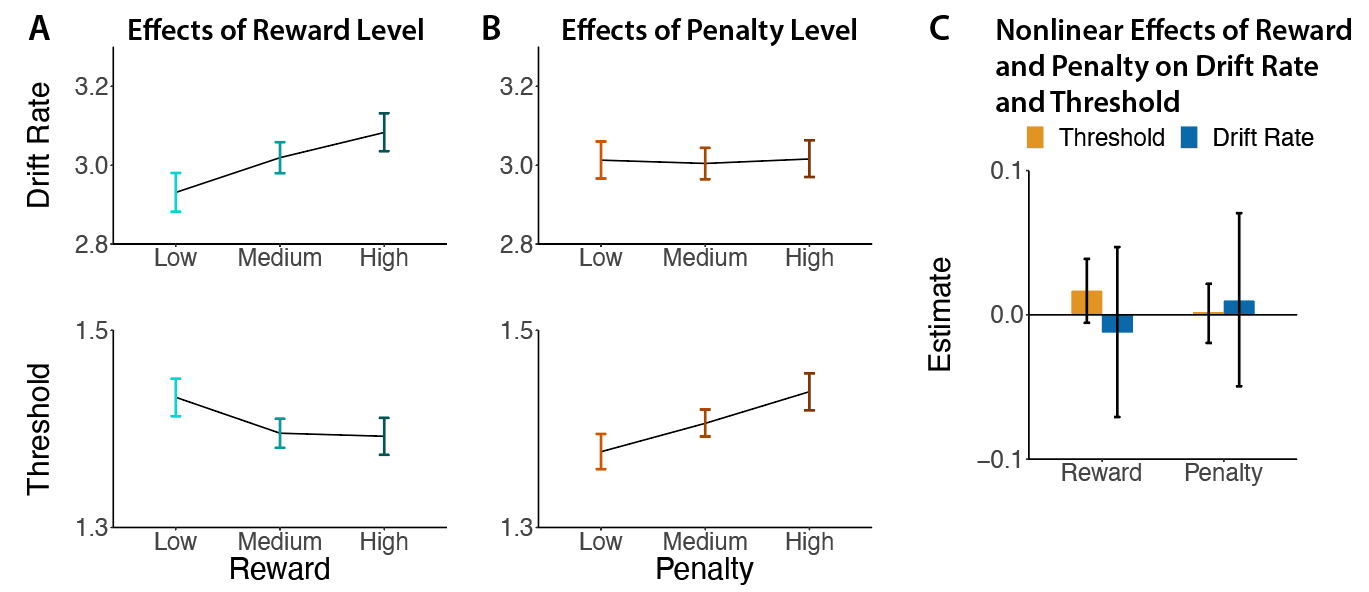

**Figure A8. Empirically observed estimates of incentive effects on DDM parameters in Study 2.** We found that **(A)** larger expected rewards led to increased drift rate whereas **(B)** larger expected penalties led to increased threshold, and that **(C)** no significant nonlinear effect of reward or penalty on DDM parameters. Error bars reflect 95% CI.

1.	Massar SAA, Pu Z, Chen C, Chee MWL. Losses motivate cognitive effort more than gains in effort-based decision making and performance. Front Hum Neurosci. 2020;14: 287.

2.	Vogel TA, Savelson ZM, Otto AR, Roy M. Forced choices reveal a trade-off between cognitive effort and physical pain. Elife. 2020;9.

3.	Soutschek A, Strobach T, Schubert T. Motivational and cognitive determinants of control during conflict processing. Cogn Emot. 2014;28: 1076–1089.

4.	Białaszek W, Marcowski P, Ostaszewski P. Physical and cognitive effort discounting across different reward magnitudes: Tests of discounting models. PLoS One. 2017;12: e0182353.

5.	Petitet P, Attaallah B, Manohar SG, Husain M. The computational cost of active information sampling before decision-making under uncertainty. Nat Hum Behav. 2021;5: 935–946.
